# Tropical land use alters functional diversity of soil food webs and leads to monopolization of the detrital energy channel

**DOI:** 10.1101/2022.02.08.479508

**Authors:** Zheng Zhou, Valentyna Krashevska, Rahayu Widyastuti, Stefan Scheu, Anton Potapov

## Abstract

Agricultural expansion is among the main threats to biodiversity and functions of tropical ecosystems. Conversion of rainforest into plantations erodes biodiversity with little-explored consequences for food-web structure and energetics of belowground communities, and associated ecosystem functions and services. We used a unique combination of approaches in stable isotope analysis and food web energetics to analyze in a comprehensive way consequences of the conversion of rainforest into plantations on the structure of and channeling of energy through soil animal food webs in Sumatra, Indonesia. Across the 23 animal groups studied, the channeling of energy shifted towards freshly-fixed plant carbon in plantations, indicating fast energy channeling as opposed to slow energy channeling in rainforests. Earthworms as major detritivores stayed unchanged in their trophic niche and monopolized the detrital pathway in plantations. Functional diversity metrics of soil food webs reflected the reduced amount of litter, tree density and species richness in plantations, providing guidelines how to improve the complexity of the structure of and channeling of energy through soil food webs. Our results highlight the strong restructuring of soil food webs with the conversion of rainforest into plantations threatening soil functioning and ecosystem stability in the long term.

## Introduction

Worldwide, land use changes the structure of ecological communities and is associated with losses in multiple ecosystem functions, which is at the core of sustainable development goals (Bommarco et al., 2013; Matson, 1997; Newbold et al., 2015). Many tropical ecosystems are affected by land-use changes, losing their biodiversity and multifunctionality (Barnes et al., 2014; Laurance, 2007). It is projected that tropical ecosystems will face even greater pressures due to land-use change in the future (Dobrovolski et al., 2011). Decreases in biodiversity and changes in trophic interactions in animal communities (Newbold et al., 2015; Tsiafouli et al., 2015; Wilkinson et al., 2021) are associated with changes in nutrient dynamics and energy fluxes (de Vries et al., 2012; McGrath et al., 2001; Potapov et al., 2020), which ultimately influence ecosystem functioning and stability (Rooney et al., 2006; Rooney and McCann, 2012). However, interrelationships between the loss of diversity and changes in energy pathways in food webs are poorly studied and this applies in particular to tropical ecosystems.

Soils harbor a large portion of terrestrial biodiversity (Guerra et al., 2021), are intimately linked with aboveground biodiversity (Bardgett and van der Putten, 2014; Hooper et al., 2000; Yang et al., 2018) and deliver vital ecosystem services (Bardgett and Wardle, 2010; de Vries et al., 2013). Energetically, 80%–90% of the carbon fixed by plants in terrestrial ecosystems enters the belowground system (Gessner et al., 2010) and is processed in soil food webs by microorganisms and invertebrate decomposers, the latter then become prey for predators (Bardgett and Wardle, 2010; Schmitz and Leroux, 2020). Shifts in resource use in the decomposer system results in asymmetries in energy fluxes through soil food web channels, which modulate the resistance and resilience of terrestrial ecosystems to perturbations (de Vries et al., 2012, 2006; Rooney et al., 2006; Rooney and McCann, 2012). Studies in temperate regions showed that more intensive land use reduces the diversity of soil organisms (Tsiafouli et al., 2015) and shifts soil food webs towards the ‘fast’ bacterial energy channel at the expense of the ‘slow’ fungal energy channel (de Vries et al., 2006), potentially undermining food web stability. However, knowledge on how the rapid land-use change in tropical regions, such as the conversion of rainforest into plantations, affects soil food web structure and energy channeling is scarce (Clough et al., 2016; Dobrovolski et al., 2011).

The present study took place in Jambi province, Sumatra, Indonesia, which is a global hot spot of biodiversity (Koh and Ghazoul, 2010; Miettinen et al., 2011), where over last 25-35 years rainforests and agroforests have been largely replaced by intensively managed plantations, mostly oil palm and rubber (Clough et al., 2016; Margono et al., 2012). Results of previous studies showed that land-use change in this region is associated with changes in soil chemistry (Ballauff et al., 2021), shifts in microbial and plant communities (Krashevska et al., 2015; Rembold et al., 2017a; Schulz et al., 2019), and reduced multitrophic biodiversity and functionality of soil animal communities (Barnes et al., 2014; Krause et al., 2021; A. M. Potapov et al., 2019a). These changes are expected to affect trophic niches of soil animals and alter both structure and energetics of soil food webs. In specific soil invertebrate groups (e.g. centipedes, springtails and mites), conversion of rainforest into plantation systems has been associated with trophic shifts towards the plant energy channel in plantations (Klarner et al., 2017; Krause et al., 2021; Susanti et al., 2021), whereas the bacterial channel was reduced, as suggested by fatty acid analysis (Susanti et al., 2019). A more complete assessment of soil food webs showed that plantations are energetically dominated by large decomposers (i.e., earthworms), but have largely reduced energy fluxes to predators (Barnes et al., 2014; A. M. Potapov et al., 2019a), however, these studies ignored potential shifts in the trophic niches of individual soil taxa with land-use change.

Progress in understanding soil food web responses to environmental changes is hampered by the chronic lack of empirical data for complex soil food webs (Brose & Scheu, 2014). Soil food webs were long reconstructed based on generic assumptions (Hunt *et al*. 1987). Insights in their structure became possible with introduction of stable isotope, molecular and biochemical methods, which showed inaccuracies in the traditional reconstructions (Bradford, 2016; Brose and Scheu, 2014; Geisen et al., 2019). Stable isotope analysis is now widely used as a first-line explorative tool in trophic ecology (Peterson and Fry, 1987; Parnell et al., 2010), allowing for an *in situ* assessment of soil food-web structure (Anton M. Potapov et al., 2019). The method is especially promising to provide insight into the structure of soil food webs in the tropics, where biology of species is poorly known. The ^13^C/^12^C and ^15^N/^14^N ratios in consumers depend on their food and can be used to explore the trophic niches of animal species and communities (D. M. Post, 2002; Pollierer et al., 2009; Anton M. Potapov et al., 2019). The ^15^N concentration is used to indicate the trophic position of species since it is enriched by about 3-4‰ per trophic level (D. M. Post, 2002; Pollierer et al., 2009; Anton M. Potapov et al., 2019); ^13^C typically is little affected by trophic transfer and thus reflects basal food resources of the trophic chain (Peterson and Fry, 1987; Anton M. Potapov et al., 2019). In soil communities, animals with high ^13^C concentration are considered to use ‘older’ carbon that have higher ^13^C values due to decomposition processes and preferential incorporation of labile plant compounds by microbes (Pollierer et al., 2009; Anton M. Potapov et al., 2019), and those with lower ^13^C values are considered to feed on freshly-fixed plant material (Fujii et al., 2021; Anton M. Potapov et al., 2019).

To assess food-web structure using stable isotope analysis, Layman et al. (2012, 2007) suggested a number of ‘isotopic metrics’, which have been widely used in aquatic ecology.

These metrics consider all species as having the same importance for food-web structure, which has a binary perspective (i.e., presence/absence of species), ignoring potential asymmetries in the magnitude of trophic interactions. However, in biological communities often only few species dominate, forming the energetic core of the food web, therefore, a non-binary perspective is important. Recently, Cucherousset and Villéger (2015) joined Layman’s metrics and the functional diversity framework (Petchey and Gaston, 2006; Villéger et al., 2008; Mouillot et al., 2013) to calculate functional diversity indices for food webs, accounting for the dominance of species. These indices include isotopic ‘richness’ which represents the volume of the trophic niche across all species, isotopic ‘evenness’ which represents the regularity of the distribution of species’ trophic niches, isotopic ‘divergence’ which reflects the dominance of species with the most extreme trophic niches, and isotopic ‘dispersion’ which reflects the balance of the species distribution in the trophic space (Cucherousset and Villéger, 2015). These isotopic indices represent basic components of functional diversity of food webs. Nevertheless, to our knowledge they have never been used to analyze soil food web characteristics, either temperate or tropical, except for one case study on oribatid mites (Krause et al., 2021).

Here, for the first time we use stable isotope analysis to comprehensively investigate changes in tropical soil food webs associated with changes in land use. We apply a functional diversity framework to stable isotope data to assess which structural dimensions of soil food webs vary most across rainforests, agroforests and intensively managed plantations of oil palm and rubber in Jambi province, Sumatra, Indonesia (Clough et al., 2016; Drescher et al., 2016). Using data on 23 high-rank taxonomic groups (orders, families), we focus on two perspectives of the functional diversity of soil food webs: a ‘community perspective’ in which we treat all groups as being equally important and an ‘energetic perspective’ in which we weight groups according to their shares in community metabolism. For both perspectives we tested the following hypotheses: (1) shifts in trophic niches are uniform across all studied animal groups through land use changes, with animals in plantations being less enriched in ^13^C than in rainforest due to stronger plant and weaker detrital energy channel; (2) functional diversity and thereby stability of soil food webs declines with land-use intensity in plantation systems reflected by reduced isotopic richness, redundancy, evenness and divergence; (3) from an ‘energetic perspective’ soil food webs are less affected by changes in land use than from a ‘community perspective’ as total energy flux changes little with conversion of rainforest into plantations, whereas biodiversity declines strongly. Lastly, we aimed at identifying the environmental factors driving changes in functional diversity of soil food webs with changes in land use from both the community and energetic perspectives.

## Materials and methods

### 1. Sampling sites

The study was conducted in the framework of the collaborative research project CRC990/EFForTS investigating ecological and socio-economic changes associated with the transformation of lowland rainforest into agricultural systems (Drescher et al., 2016). Four land-use systems, rainforest, jungle rubber, rubber plantations and oil palm plantations were investigated in two regions, i.e. Harapan and Bukit Duabelas (Drescher et al., 2016). Jungle rubber sites were established by planting rubber trees (*Hevea brasiliensis*) into selectively logged rainforest and contain rainforest tree species. Jungle rubber sites represent low intensive land-use systems, lacking fertilizer input as well as herbicide application; the age of rubber trees varied between 15–40 years (Kotowska et al., 2015). Rubber and oil palm (*Elaeis guineensis*) monocultures represent high land-use intensity plantation systems managed by the addition of fertilizers as well as herbicides (Drescher et al., 2016). Each land-use system was replicated four times in each landscape, resulting in a total of 32 sites; for more details see Drescher *et al*. (2016).

### 2. Sampling, extraction and classification of soil fauna

Soil animals were sampled at each of 32 study sites during October and November 2013. Soil samples measuring 16 cm x 16 cm and including the litter layer and 0 – 5 cm of the mineral soil were taken in three 5 m × 5 m subplots within each of 50 m × 50 m plots established at each study site, resulting in a total of 96 samples. The samples were transported to the laboratory and animals were extracted by heat (Kempson et al., 1963) until the substrate was completely dry (6–8 days). Until further analysis, species were stored in 70% ethanol. For calibration of the animal stable isotope values, we used mixed litter samples that were taken from each site and analyzed in a previous study (Klarner et al., 2017).

Animals were classified into 23 high-rank taxonomic groups (orders, families). For stable isotope analysis we used this “group-level approach” to allow generalization across the whole soil animal food web. High-rank animal taxa in soil are generally consistent in their isotopic niches and reflect the trophic niches of species in most taxa (A. M. Potapov et al., 2019b). We further classified taxonomic groups into five major functional groups according to their trophic guild and body size class (A. M. Potapov et al., 2021, 2019a): herbivores including e.g., Hemiptera and Orthoptera, microdecomposers including e.g., Oribatida and Collembola, macrodecomposers including e.g., Annelida and Diplopoda, micropredators including e.g., Diplura and Mesotigmata, macropredators including e.g., Araneae and Chilopoda, and groups with mixed feeding habits including e.g., Diptera and Coleoptera.

### 3. Stable isotope analysis

To cover the entire community, for each sampling site we analyzed all taxa for which we were able to collect enough biomass for stable isotope analysis and which were represented by more than two individuals (Table S1). We analyzed a minimum of 3 and a maximum of 15 individuals for each taxonomic group for each site as a single mixed sample to cover the species- and individual-level isotopic variation. We mixed individuals from different subplots whenever possible to cover spatial variation in stable isotope values. Animals from the litter and soil layer were analyzed separately, but were merged for data analysis since stable isotope values did not differ significantly between layers. Animal samples were dried at 60°C for 24 h before stable isotope analysis, weighed and wrapped into tin capsules; sample weights varied between 0.01 and 1.00 mg. For small-sized animal groups we used bulk individuals, for large-sized animal groups we used body parts dominated by muscle tissue (e.g., legs) from different individuals and pooled them (Tsurikov et al., 2015). In total, 626 samples of 23 taxonomic groups were analyzed across 32 sites. For Collembola, Oribatida and Chilopoda we additionally used stable isotope data collected at species level (Klarner et al., 2017; Krause et al., 2019; Susanti et al., 2021) to calculate a single average value for each group at each site. The number of analyzed taxonomic groups varied between 6 and 17 per site (i.e., per one soil food web) and was on average 12.3.

Animal samples were analyzed using a coupled system of an elemental analyzer (NA 1500, Carlo Erba, Milan, Italy) and a mass spectrometer (MAT 251, Finnigan, Bremen, Germany) adopted for the analysis of small sample sizes (Langel and Dyckmans 2014). Ratios of the heavy isotope to the light isotope (^13^C/^12^C,^15^N/^14^N, denoted as R) were expressed in parts per thousand relatives to the standards using the delta notation with δ^13^C or δ^15^N = (R_sample_/R_standard_ - 1) × 1000 (‰). Vienna PD Belemnite and atmospheric nitrogen were used as standard for ^13^C and ^15^N respectively. Acetanilid was used for internal calibration.

Environmental parameters of the study sites were used as given in Potapov *et al*. (2020), Krashevska *et al*. (2017) and Rembold *et al*. (2017a).

### 4. Statistical analysis

The stable isotope compositions of animals were calibrated to that of the local leaf litter. Calibrated δ^13^C and δ^15^N values were calculated as the difference between the plot-specific litter δ^13^C and δ^15^N values and the δ^13^C and δ^15^N values of each group, and given as Δ^13^C and Δ^15^N values, respectively. Statistical analyzes were done in R v 4.0 (R Core Team, 2020) with R studio interface (RStudio Team, 2020).

To characterize the trophic structure of soil animal communities we calculated isotopic metrics as given in Cucherousset and Villéger (2015). One-dimensional metrics describe the isotopic parameters of the communities based on Δ^15^N or Δ^13^C values. Multidimensional metrics combine both Δ^13^C and Δ^15^N values, and join the ones from Layman et al. (2007), with functional diversity framework (Villéger et al. (2008); Laliberté and Legendre (2010). The Δ^13^C and Δ^15^N values were scaled between 0 and 1 based on maximum and minimum across all communities to ensure equal contribution of two isotopes prior to calculation of multidimensional metrics. Multidimensional metrics were calculated from two perspectives: (1) a ‘community perspective’, assuming all taxonomic groups being equally important, i.e. *unweighted metrics*, and (2) an ‘energetic perspective’, assuming that groups that have higher contribution to total community metabolism are also more functionally important, i.e. metrics were *weighted by community metabolism*. We used metabolism instead of biomass because it better reflects the contribution of organisms to energy processing and thus their importance in the food web (Barnes et al., 2018; Brown et al., 2004). Community metabolism for each group at each plot was taken from Potapov *et al*. (2019a); it was based on length and width measurements of all individuals and using body size to body mass ratios and group-specific allometric regressions to calculate metabolic rates (Ehnes et al., 2011). Individual metabolic rates were then summed up for groups to estimate contribution of each taxonomic group to the total community metabolism per plot (Table S1).

Overall, thirteen isotopic metrics were calculated for each of 32 communities (i.e., sampling plots). One-dimensional metrics included average position, range, minimum and maximum. The unweighted and metabolism-weighted *average position* of communities (mean isotopic value across groups) represent mean community-level isotopic trait values. The *isotopic range* represents the difference between *minimum* and *maximum* values of both Δ^13^C and Δ^15^N. Range, minimum and maximum could not be weighted and are given unweighted. Multidimensional metrics included isotopic divergence, isotopic dispersion, isotopic evenness, isotopic uniqueness and isotopic richness, which were calculated as both unweighted and metabolism-weighed. *Isotopic divergence* represents the distance between all species and the center of the convex hull area. Isotopic divergence values close to 0 indicate that groups with extreme stable isotope values are rare (community divergence) or contribute little to the community metabolism (energetic divergence), whereas isotopic divergence values close to 1 indicate that there are many groups with extreme stable isotope values (community divergence) or they contribute considerably to the community metabolism (energetic divergence). *Isotopic dispersion* combines convex hull area with isotopic divergence values and can be interpreted as scaled multidimensional variance. Isotopic dispersion approaches 1 when species with contrasting stable isotope values have similar abundance, which is a more functionally diverse and balanced system, whereas it approaches 0 when most groups (community dispersion) or community metabolism (energetic dispersion) are concentrated near the ‘center of gravity’ of the community in stable isotope space. *Isotopic evenness* quantifies the distribution of groups or metabolism in stable isotope space. Isotopic evenness values close to 1 indicate that the isotope values of the groups/metabolism are evenly distributed, while values close to 0 indicate that the groups/metabolism cluster together. *Isotopic uniqueness* reflects the closeness of stable isotope values of the studied groups/metabolism within the community, which is defined as the inverse of the average isotopic redundancy. Finally, *isotopic richness* is the volume occupied by all groups in isotopic space (convex hull area in two-dimensional isotopic space) and reflects functional richness of the food web; it is the only multidimensional metric that cannot be weighted since it considers the total isotopic space (Mason et al., 2005; Villéger et al., 2008).

To assess differences in food-web structure among land-use systems we used a set of analyzes of variance (*aov* function) with the Δ^13^C and Δ^15^N values of each taxonomic group, one-dimensional isotopic metrics, and multidimensional community and energetic isotopic metrics as response variables, and land-use system (rainforest, jungle rubber, rubber, oil palm) and landscape (Harapan or Bukit Duabelas) as factors (total n = 32, 8 plots as replicates per land-use system). Pairwise comparisons of means among land-use systems were done using post-hoc *HSD.test* function from the package *agricolae* (Mendiburu, 2020) following analyzes of variance. Differences in Δ^13^C and Δ^15^N values between rainforest and other land-use systems for each taxonomic group were analyzed with Student’s *t.test* function in R. Results were visualized using the *ggplot2* package (Wickham, 2016).

To assess effect size of land use on all food-web metrics combined, we used analysis of similarities based on community and energetic metrics with land use as the grouping variable (*anosim* in package *vegan*). Besides, we used multivariate analyzes of variance (MANOVAs) to inspect the effects of environmental factors on community and energetic metrics, and additionally explored pairwise correlations between environmental factors and food-web metrics using Spearman’s correlation from the package *agricolae* (Mendiburu, 2020).

Finally, a structural equation model (SEM) based on generalized least squares was constructed to provide insight into how land use affected soil food webs from both community and energetic perspectives. The analysis was performed with the *lavaan* package in R (Rosseel, 2012). The model included tree, understory and soil properties selected according to permutation tests based on R^2^ which were used to quantify the land-use effects (*ordiR2stepin* package *vegan*), and before permutation tests, the environmental factors were filtered based on the MANOVAs and Spearman’s correlation. The final model included soil pH, tree density and tree richness as the three most important variables that represented direct land-use effects (i.e., logging and liming applications; Drescher *et al*. 2016). Furthermore, we included litter amount, understory density and earthworm metabolism as the three mediators that are affected by changes in tree density, tree richness and pH, and have strong impacts on tropical soil invertebrate communities (Darras et al., 2019; A. M. Potapov et al., 2019a). Food-web metrics (i.e., isotopic divergence, dispersion, uniqueness, evenness and average position) were first combined using Principal Component Analysis (*prcomp* function) and the PC1 was used as the response variable in SEM; PC1 explaned 66% and 42% of the variance for community and energetic metrics, respectively. To determine the goodness of fit of the model we used χ^2^-test associated P-value ≥ 0.05, the comparative fit index (CFI) > 0.95, the root-mean-square error of approximation (RMSEA) and the standardized root-mean-square residual (SRMR) with values ≤ 0.05 (Schermelleh-Engel et al., 2003). Our SEM adequately described the data (χ^2^ = 6.23, *p* = 0.40, df = 5, CFI = 0.98, RMSEA = 0.04, SRMR = 0.05).

## Results

### 1. Isotopic shifts in individual animal taxa

If averaged across rainforest sites, the mean Δ^13^C values of taxonomic groups covered the range of 3.0‰, from 3.4‰ (Coleoptera and Oribatida) to 6.4‰ (Orthoptera and Pauropoda). The respective Δ^15^N values covered the range of 14.2‰, from −5‰ (Pauropoda) to 9.2‰ (Diplura; Fig. 1a). The Δ^13^C values of all groups varied across the four land-use systems, but Chilopoda, Diplura and Annelida were typically most enriched in ^13^C and Coleoptera was consistently among the most depleted in ^13^C among all groups. Diplura, Preudosorpiones, Chilopoda and Isopoda had the highest Δ^15^N values among all groups across the four land-use systems. The group with lowest Δ^15^N values was Pauropoda in rainforest and jungle rubber and Protura in rubber and oil palm plantations. Overall, micropredators, i.e., Diplura and Preudosorpiones, had 2-3‰ higher Δ^15^N values than macropredators, i.e., Chilopoda, Formicidae and Araneae (Fig. 1). The share of Annelida (earthworms) in community metabolism was 15.4% in rainforest, but represented more than 75% in jungle rubber, rubber and oil palm plantations (Fig. S1).

**Figure 1.**
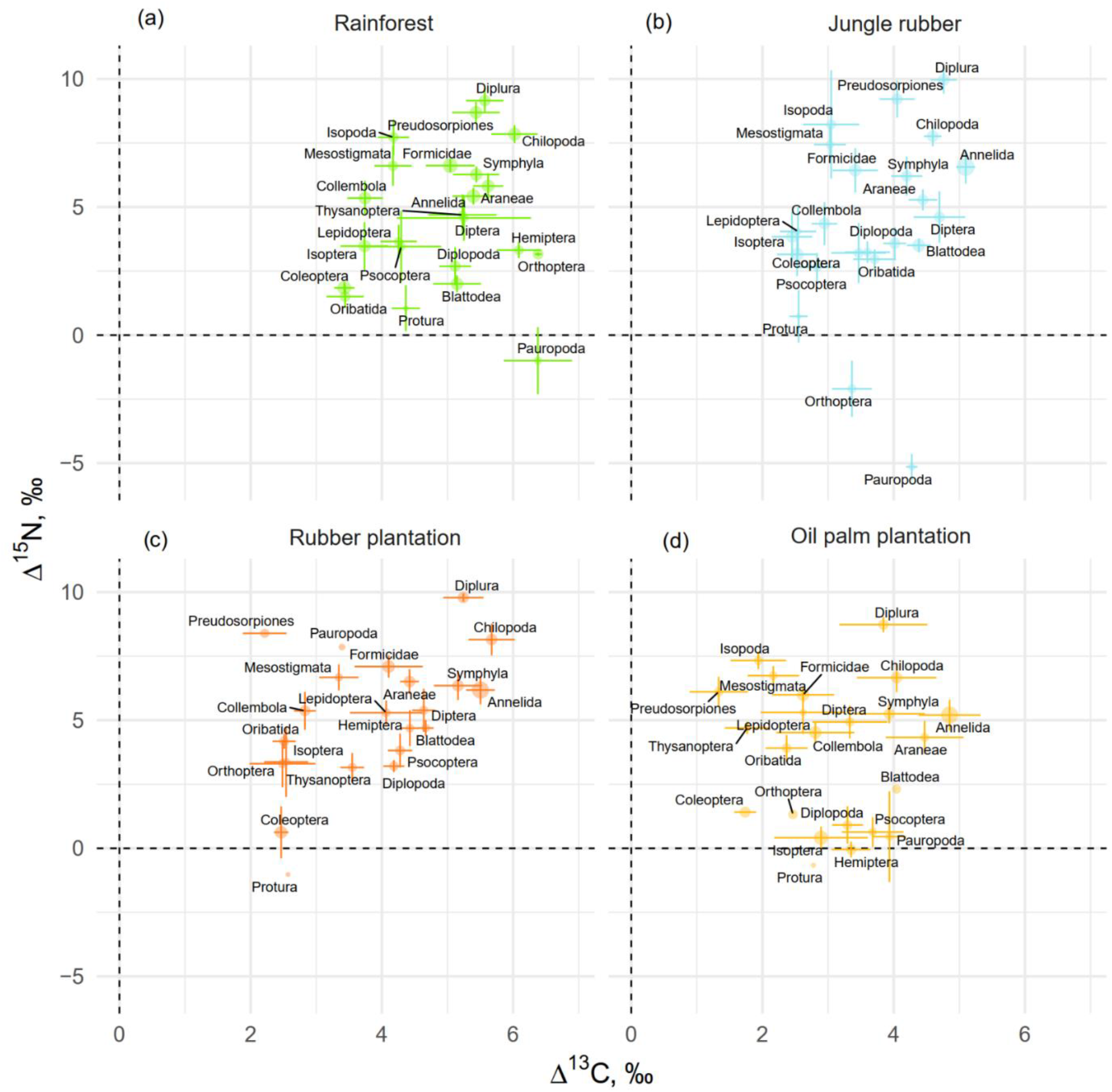
Mean litter-calibrated Δ^13^C and Δ^15^N values of soil animal taxa in rainforest (a), jungle rubber (b), rubber (c) and oil palm plantations (d). Error bars represent standard errors across sampling plots (n = 1-8 per land-use system). Size of the points is scaled to the total share of the taxonomic group in the community metabolism in the corresponding land-use system (metabolism was log_10_-transformed to show trends in rarer groups).

The Δ^13^C values were significantly higher in rainforest than in other land-use systems in Coleoptera, Diplopoda, Hemiptera, Orthoptera, Pauropoda, Protura, Preudosorpiones and Thysanoptera (Fig. 2, Fig. S2). Chilopoda, Diplura, Formicidae, Isopoda, Mesostigmata and Symphyla were significantly more enriched in ^13^C in rainforest than in oil palm, but not significantly different from those in jungle rubber and rubber plantations. In general, most groups in rainforest were higher in Δ^13^C by 1-3‰ than in the other land-use systems, but this shift was only significant for two out of six macrodecomposer groups. Annelida, which accounted for much of the community metabolism in each of the land-use systems, had similar Δ^13^C values across land-use systems.

**Figure 2.**
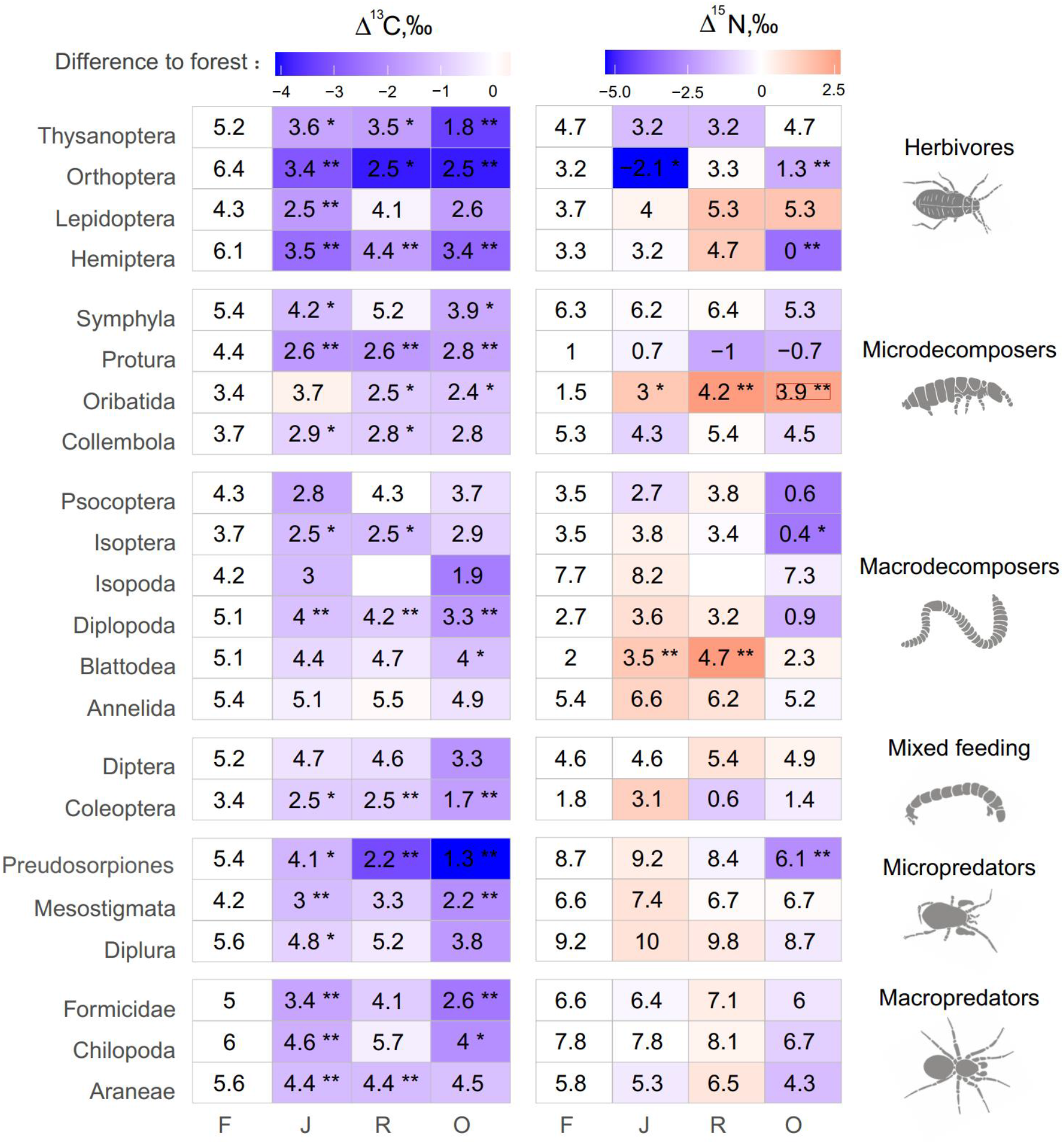
Average Δ^13^C and Δ^15^N values of taxonomic groups in rainforest (F), jungle rubber (J), rubber (R) and oil palm plantations (O). Numbers show means, asterisks indicate significant differences between the mean value in the corresponding land-use system and in rainforest (Student’s *t*-test \**p* <0.05, \*\**p*<0.01). Color represents the direction (red – increase, blue - decrease) and magnitude (brighter color indicate stronger change) of the difference between rainforest and other land-use systems.

The Δ^15^N values were by 1.5-2.5‰ lower in rainforest than in the other land-use systems in Oribatida and Blattodea (except for similar Δ^15^N values in oil palm and rainforest for Blattodea). By contrast, Δ^15^N values of Hemiptera, Orthoptera, Isoptera and Preudosorpiones were lower in oil palm than in rainforest, whereas in jungle rubber this was only true for Orthoptera (Fig. 2, Fig. S3).

### 2. One-dimensional isotopic metrics

One-dimensional isotopic metrics described the overall range and average Δ^13^C and Δ^15^N values of each community. The maxima of Δ^13^C values were by 1-2‰ higher in forest than in jungle rubber and oil palm plantations, but minima and the overall range did not differ significantly (Fig. 3). The unweighted average Δ^13^C values of communities were by 1-2‰ higher in rainforest than in the other land-use systems and were also higher in rubber than in oil palm plantations. However, the energetic average positions did not differ significantly due to similar Δ^13^C values of Annelida (dominant invertebrate group) across land-use systems (Fig. 2e, Fig. S1).

**Figure 3.**
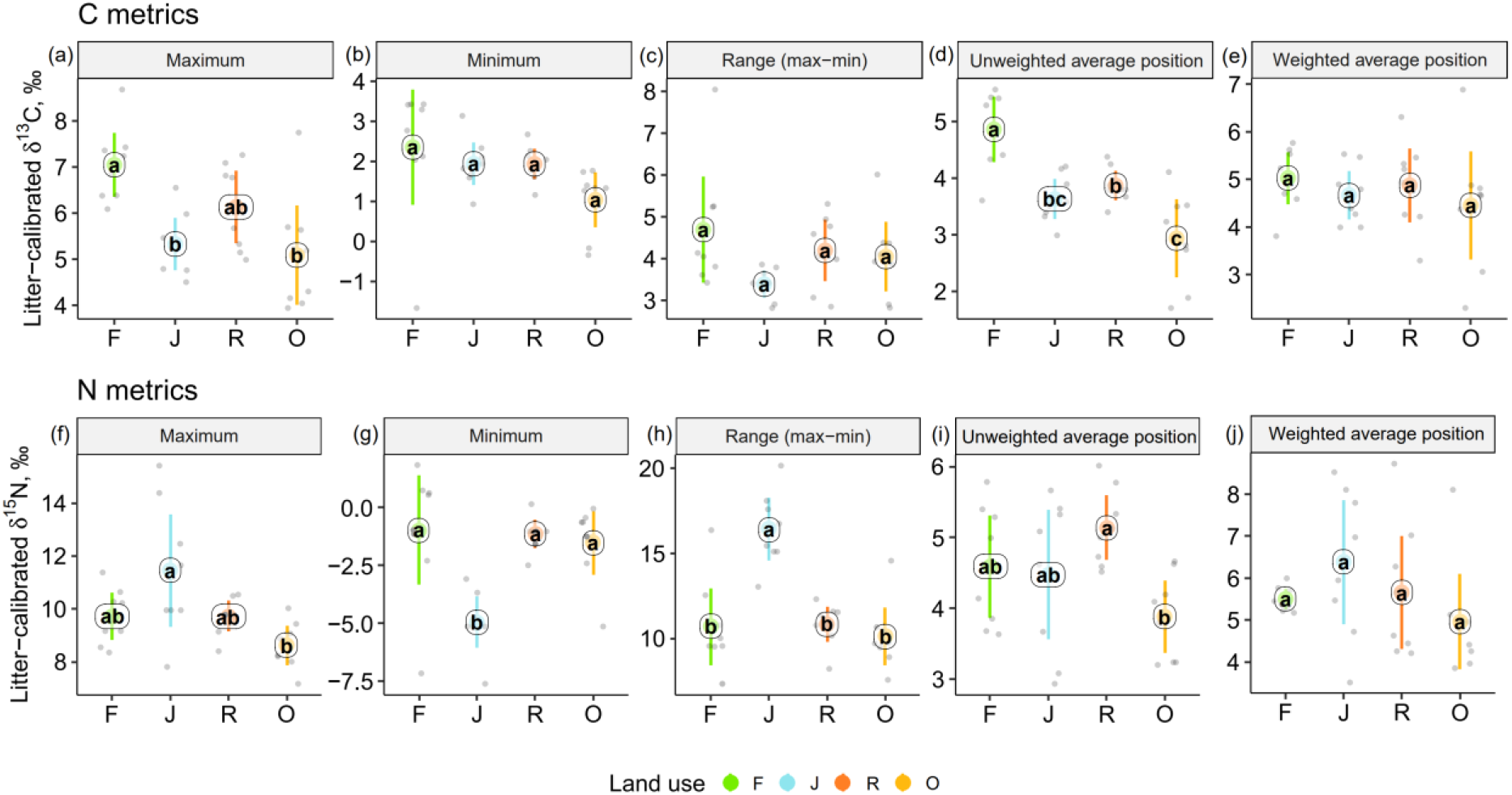
One-dimensional metrics for Δ^13^C (upper panel) and Δ^15^N values (lower panel) of communities in rainforest (F, green), jungle rubber (J, blue), rubber (R, red) and oil palm plantations (O, yellow). Each point represents one community (n = 8 per land-use system). For the calculation of the weighted average values, species were weighted according to their contribution to the total community metabolism per plot. Means, sharing the same letter within each pane are not significantly different (Tukey’s HSD test following ANOVA, *p* > 0.05).

Extreme values of Δ^15^N were most pronounced in jungle rubber, both maximum and minimum, resulting in the largest range in Δ^15^N values (16.5‰ among the four land-use systems. In the other land-use systems, maxima, minima and ranges of Δ^15^N values were similar. The unweighted average Δ^15^N values of communities were lowest in oil palm, being significantly lower than in rubber (Fig. 3i). By contrast, the energetic average Δ^15^N values did not differ significantly.

### 3. Multidimensional isotopic metrics

Among the unweighted metrics (i.e., community metrics), isotopic dispersion was significantly higher in oil palm than in each of the other land-use systems; isotopic divergence and uniqueness were significantly higher in oil palm than in jungle rubber; isotopic evenness was significantly lower in jungle rubber than in each of the other land-use systems; only isotopic richness showed no significant differences between land-use systems, but in trend the two monoculture systems had lower values than in rainforest and jungle rubber. For detailed information on the plot-level metrics values see Appendix (Figs S4 – S36).

By contrast, the weighted multidimensional metrics (i.e., energetic metrics) did not differ among land-use systems for isotopic dispersion, isotopic evenness, isotopic richness and isotopic uniqueness (Fig. 4). Only isotopic divergence was significantly lower in rainforest than in the other land-use systems, showing an opposite trend to isotopic dispersion. For detailed information on plot level metrics values see Appendix (Figs S39 – S68).

**Figure 4.**
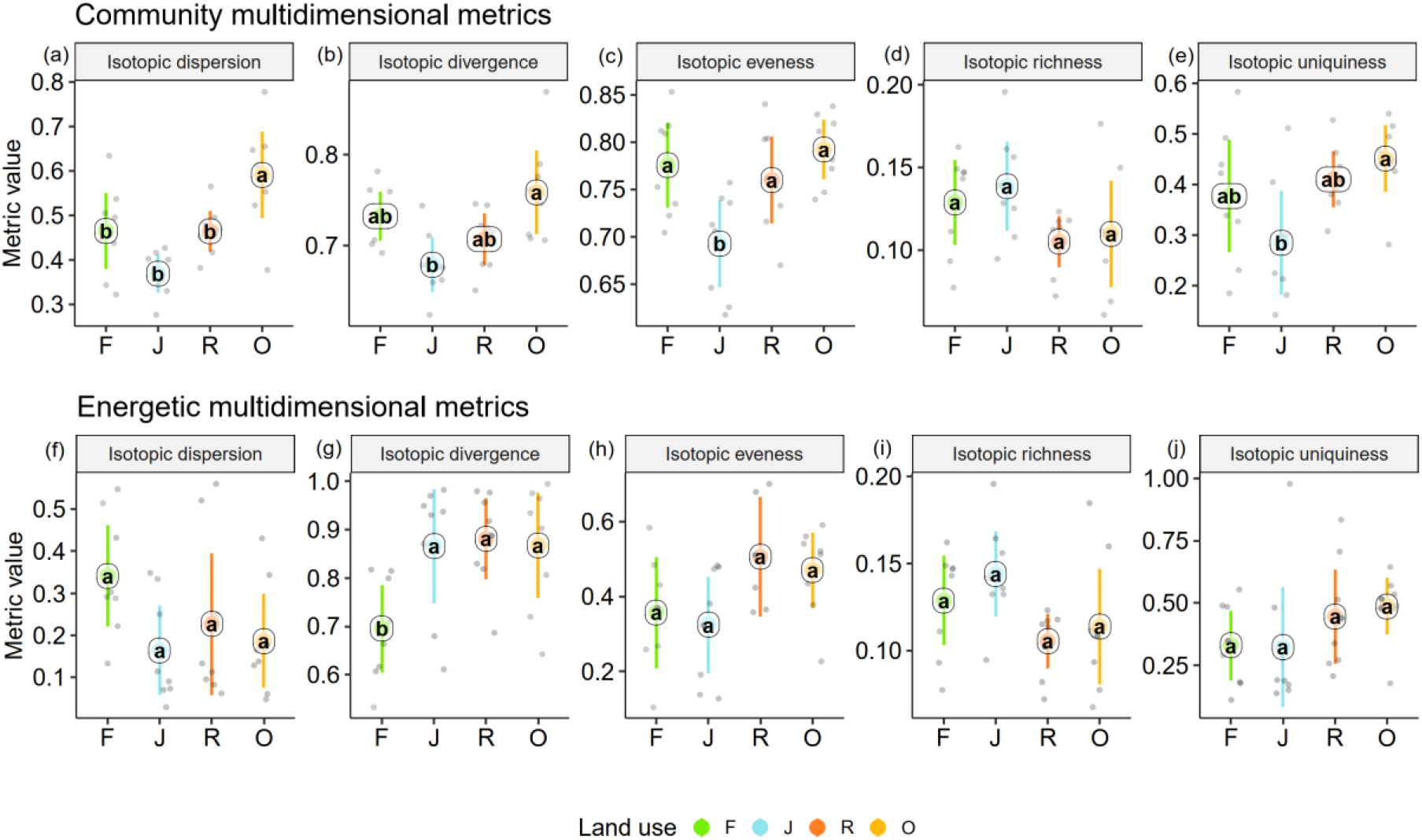
Multidimensional isotopic metrics of soil animal communities in rainforest (F, green), jungle rubber (J, blue), rubber (R, red) and oil palm plantations (O, yellow). Community (upper panel) and energetic metrics (lower panel) are shown. Each point represents one community (n = 8 per land-use system). Means sharing the same letter within each pane are not significantly different (Tukey’s HSD test following ANOVA, *p* < 0.05).

### 4. Environmental effects on functional diversity of soil food webs

As indicated by multivariate analysis, community metrics differed strongly between the four land-use systems (anosim R = 0.404, *p* < 0.001), whereas differences for the energetic metrics were less pronounced (anosim R = 0.138, *p* = 0.014). Among all tested environmental factors, soil pH, tree species richness, litter amount and understory density had the strongest correlations with both community metrics (*p* = 0.003, *p* = 0.037, *p* < 0.001, *p* < 0.001, respectively) and energetic metrics (*p* = 0.004, *p* = 0.002, *p* = 0.003 *p* = 0.002, respectively).

These variables were subsequently selected for the SEM analysis (see methods). SEM indicated that the changes in the community metrics (PC1_unweighted_) were induced directly by tree properties and litter amount (tree density: *p* < 0.05, effect size = 0.72; tree species richness: *p* < 0.001, effect size = −0.83; litter amount: *p* < 0.001, effect size = 0.71), while changes in the energetic metrics (PC1_weighted_) were indirectly driven by soil pH via increased metabolism of earthworms (*p* < 0.05, effect size = −0.36; Fig 6).

**Figure 5.**
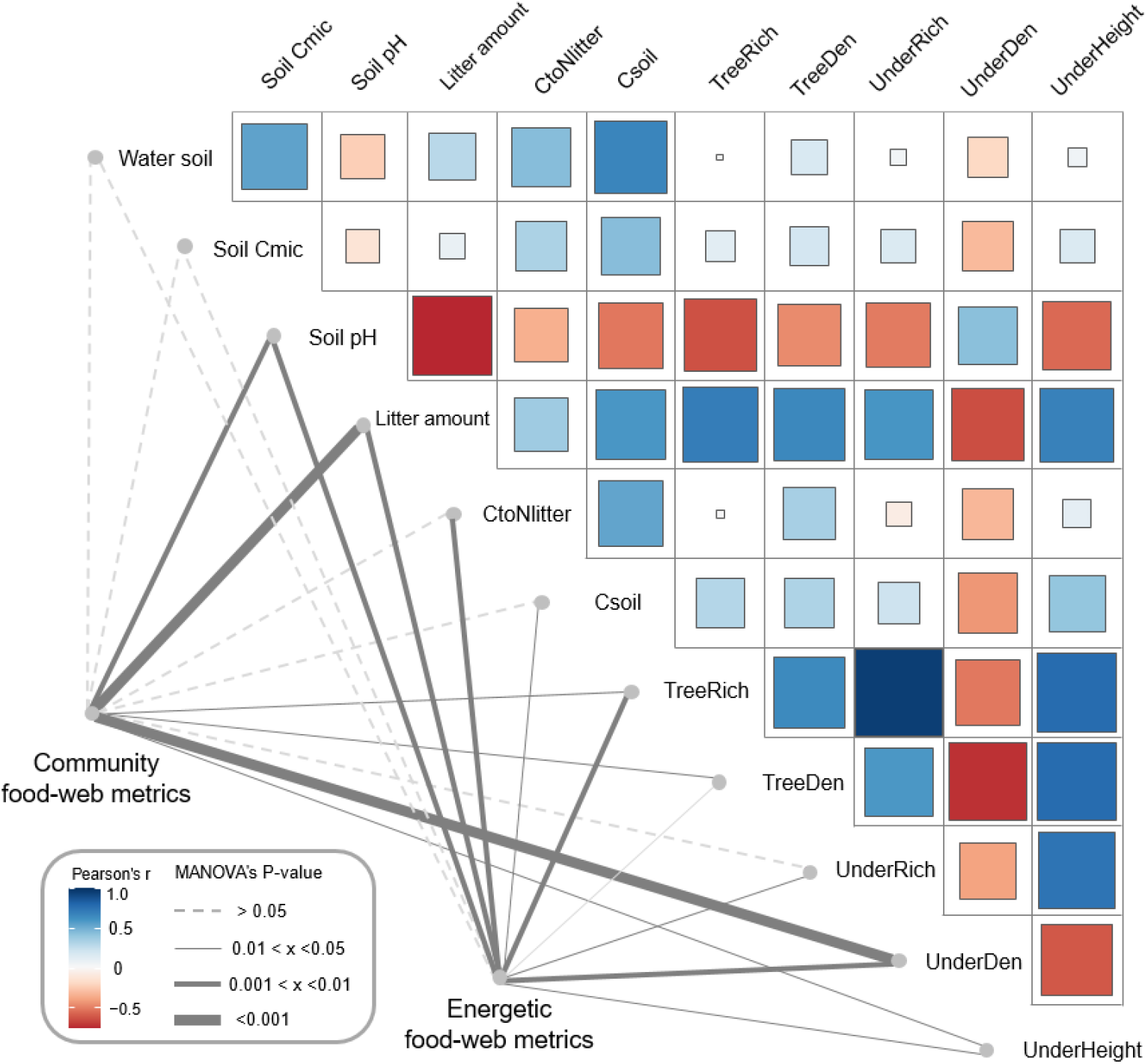
Environmental drivers of community and energetic soil food-web metrics. Community and energetic food-web metrics were related to environmental factors using MANOVA; the thickness of connection lines shows statistical significance, dashed line for *p* > 0.05. Pairwise Spearman’s correlations among environmental factors are shown with a tile chart (blue – negative, red – positive). The vegetation parameters included tree species richness (TreeRich), tree density (TreeDen), understory species richness (UnderRich), understory density (UnderDen), and average understory height (UnderHeight). Parameters of litter and soil include soil pH, litter amount, soil carbon concentration (Csoil), carbon to nitrogen ratio of litter (CtoNlitter), soil microbial biomass C (Soil Cmic) and soil humidity (Water soil).

**Figure 6.**
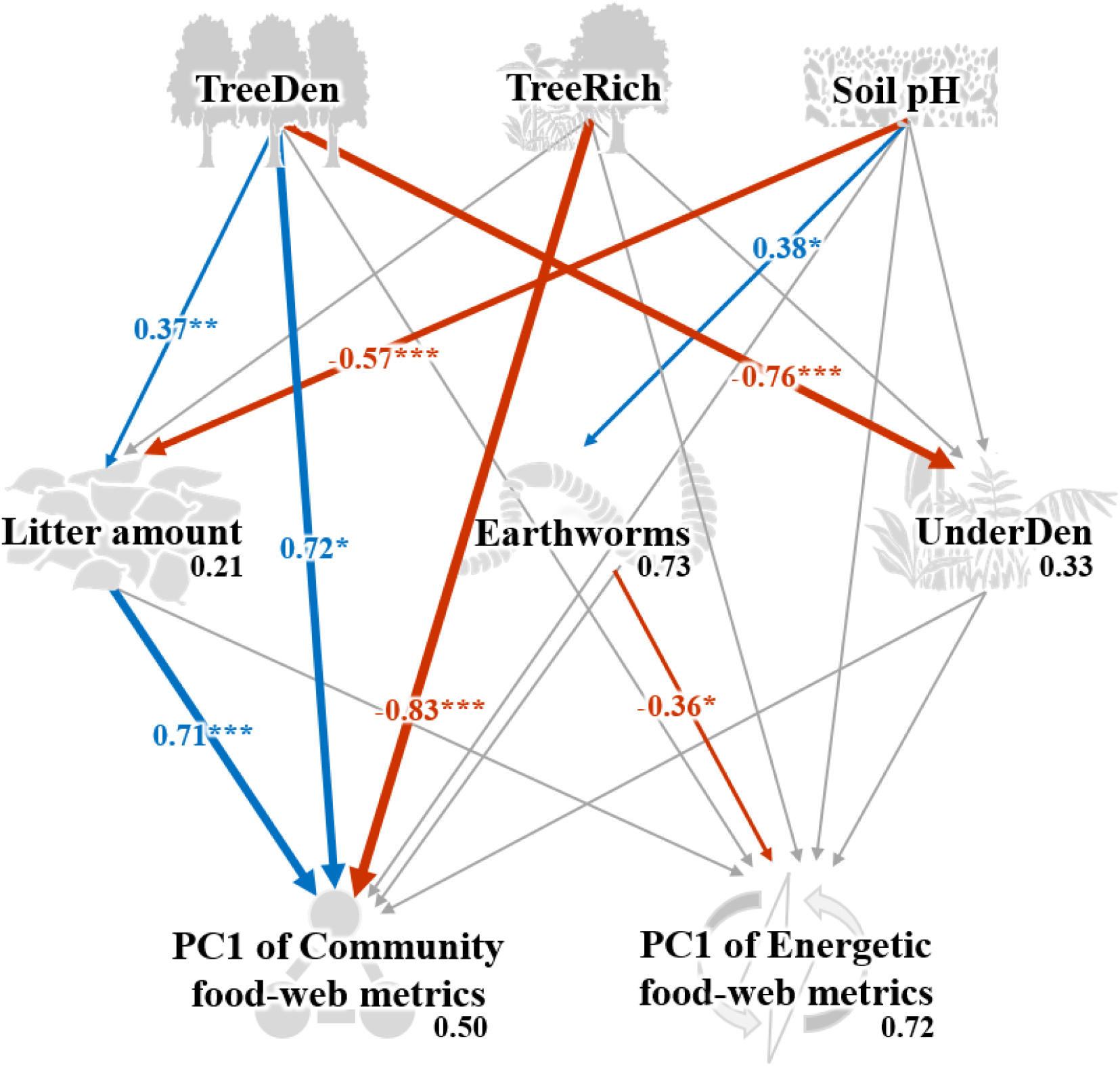
Structural equation model on the effects of environmental change on food-web metrics. Numbers adjacent to arrows are standardized path coefficients that show effect sizes and directions (blue – positive, red – negative) of the relationship, arrow width is proportional to the strength of path coefficients. Grey arrows represent paths that were not significant; \**p* < 0.05, \*\**p* < 0.01, and *\*\*\*p* < 0.001. Numbers above every response variable in the model denotes the proportion of variance explained. For abbreviations see Figure 5.

## Discussion

We used stable isotope data of 23 high-rank animal taxa to comprehensively assess changes in functional diversity of soil food webs under tropical land-use change. We found shifts in basal resource use for most of taxonomic groups in plantations and responses of food-web diversity metrics to land use that were more pronounced for community than for energetic metrics. In agreement to our first hypothesis, ^13^C values of animal taxa and communities were more enriched in rainforest than in plantations, but this shift vanished if the average Δ^13^C values were weighted by metabolism. Jungle rubber had the largest range of Δ^15^N values among all land-use systems, which suggests the longest food-chain in this system. Refuting our second hypothesis, when considering all taxonomic groups being equally important (‘community perspective’), soil food webs in oil palm had a significantly higher community dispersion than in the other land-use systems and in trend also had a higher community divergence and uniqueness (i.e., a lower redundancy). By contrast, most energetic isotopic metrics (‘energetic perspective’) varied less between the four land-use systems. Conform to our third hypothesis, community isotopic metrics were more sensitive to changes in land use than energetic isotopic metrics, suggesting soil food webs are more sensitive to land-use change from a community than from an energetic perspective. Further, community metrics of soil food webs were influenced directly by land use (tree properties and litter amount), whereas energetic metrics were influenced indirectly via pH-induced changes in earthworm abundance.

### 1. The structure of tropical soil food webs

Our study is among the first comprehensive assessments of tropical soil food webs based on stable isotope analysis. Collembola and Isopoda showed a much higher ^15^N enrichment than e.g., Oribatida, but all three groups occupy similar trophic positions in temperate forests and predominantly function as decomposers (Anton M. Potapov et al., 2019). Protura in temperate forests are enriched in ^15^N and feed on ectomycorrhizal fungi (Bluhm et al., 2019), whereas in the studied tropical forests, Protura had the lowest Δ^15^N values among all groups, suggesting that they feed on saprotrophic rather than mycorrhizal fungi (Bluhm et al., 2019). The low Δ^15^N and high Δ^13^C values of Pauropoda, first even reported for this group, indicate that they function as decomposers by feeding on saprotrophic microorganisms (Tiunov et al., 2015), confirming earlier suggestions (Starling, 1944). Low Δ^15^N values in a number of taxa may be associated with feeding on algae (Potapov et al., 2018), shown to be important for mesofauna in tropical soil food webs (Susanti et al., 2019). Unexpectedly, micropredators (e.g., Diplura and Pseudoscorpiones) had higher trophic positions (Δ^15^N values) than macropredators (e.g., Araneae and Formicidae) across all land-use systems, and Diplura had the highest Δ^15^N values among all taxa studied. Diplura were represented mostly by predatory Japygidae, which may hunt springtails, mites and other small invertebrates (Sendra et al., 2021). The higher trophic position of small-sized predators suggest that they form part of a different energy channel than macropredators. In fact, the micro-food web in soil has been shown to be based mainly on microbial resources channeled to higher trophic levels by microarthropod predators, whereas the macro-food web is based more on litter and detritus consumed by macrofauna taxa with the energy channeled to higher trophic levels by macroarthropod predators (A. M. Potapov et al., 2021). This implies more trophic transactions in the micro-food web (Pollierer et al., 2009; Steffan et al., 2015) explaining the higher trophic position of micro-than macroarthropod predators.

### 2. Shifts in trophic positions of soil invertebrates and energy channels in soil food webs

In agreement with our first hypothesis, Δ^13^C values of most of the studied soil animal taxa were higher in rainforest than in plantations. The ^13^C concentration in dead plant material is increasing during decomposition compared to fresh leaf litter (Ågren et al., 1996; Boström et al., 2007; Anton M. Potapov et al., 2019), and high Δ^13^C values in soil fauna in forest is likely indicates feeding on saprotrophic fungi and bacteria that assimilate predominantly labile ^13^C-enriched plant compounds (Hyodo, 2015; Pollierer et al., 2009; Potapov et al., 2013). Vascular and non-vascular plants have generally lower Δ^13^C values than saprotrophic microorganisms and animals (Hyodo et al., 2010; Anton M. Potapov et al., 2019), therefore, the high Δ^13^C values in soil invertebrates in rainforest point to a more pronounced detritus-based ‘brown’ food web relying heavily on saprotrophic fungi and bacteria based on litter material. Among the plantations, the unweighted average Δ^13^C values were lowest in oil palm suggesting a shift towards a more plant-based ‘green’ food web relying more heavily on the consumption of living plant tissue (Fujii et al., 2021), which has been previously shown for Chilopoda, Oribatida, Collembola and Pseudoscorpiones on the same study sites (Klarner et al., 2017; Krause et al., 2019; Liebke et al., 2021; Susanti et al., 2021). Results of the study of Susanti *et al*. (2019) at our study sites further support the conclusion of a more pronounced plant- and reduced detritus-based energy channel in soil food webs of plantations compared to rainforest using fatty acids as trophic biomarkers. Compared to rainforest the herb layer is much more developed in plantations due to more open canopy and coverage by weeds (Rembold et al., 2017a), presumably providing high quality resources for plant and litter feeding soil animals. By contrast, litter in rainforest is high in lignin and therefore of low food quality (Krashevska et al., 2018), increasing the use of saprotrophic microorganisms rather than litter by detritivores (Illig et al., 2005).

In soil food-web models the bacterial energy channel typically is considered to be ‘fast’, while the fungal energy channel is considered to be ‘ slow’ due to differences in the turnover rates of bacteria and their consumers, and fungi and their consumers, respectively (Coleman et al., 1983; Moore et al., 2005). Generally, the fast energy channel is defined by fast-growing populations with short turnover rates (Rooney et al., 2006). Based on this concept, animals feeding on living plants or fresh litter contribute to fast energy channeling, whereas animals feeding on recalcitrant detritus (being slowly decomposed belowground) are associated with slow energy channeling. This perspective complements the green (plant) and brown (detrital) energy channels in soil food webs. Since ^13^C accumulates in plant-derived materials with microbial decomposition (Fujii et al., 2021; Pollierer et al., 2009; Potapov et al., 2013), Δ^13^C values in soil animals may indicate slow versus fast carbon cycling in soil. Therefore, the shift towards plant–based energy channeling in food webs of plantations may reflect accelerated carbon cycling at the ecosystem level, contributing to carbon losses of plantations (Guillaume et al., 2018) and thereby potentially compromising ecosystem long-term stability (McCann *et al*. 1998; Rooney & McCann 2012).

Contrasting the community perspective, energetic Δ^13^C values of communities did not vary significantly among land-use systems, which was due to similar Δ^13^C values of Annelida (earthworms) and some other macro-decomposers across land-use systems (Fig.2, Fig. S1). Earthworms had the highest share in community metabolism among detritivores in plantations suggesting that they predominate animal-mediated decomposition and organic matter transformation processes (A. M. Potapov et al., 2019a). Notably, earthworms were among the most ^13^C-enriched animal groups in jungle rubber, rubber and oil palm plantations, but their Δ^13^C values were similar to other soil animal groups in rainforest. earthworms feed on detritus and microorganisms and are able to efficiently use fresh litter carbon, but also ‘old’ microbially-processed carbon (e.g., soil organic matter) (Blouin et al., 2013; Hyodo et al., 2012; Scheu and Falca, 2000) reflected in high Δ^13^C values (Pollierer et al., 2009). Land-use change in tropical lowland landscapes is associated with losing biodiversity and biomass of litter arthropods (Barnes et al., 2014), but the negative effect of biodiversity loss on soil functions may be at least in part counteracted by earthworms that monopolize the detrital channel in plantations. Thereby, earthworms also may counteract the destabilization of the system through sequestration of carbon in their large body and thus strengthening ‘slow’ energy channeling (Rooney and McCann, 2012; Schwarzmüller et al., 2015). Earthworms contribute only about 15.4% to community metabolism in rainforest, leaving vacated trophic space for other groups that are vanish or reduced under land-use change. Combined, high Δ^13^C values and shift in dominance of detritivore taxa suggest that the detrital energy channel in rainforest is diversified and comprised a wider range of consumer groups than in plantations, whereas in plantations it comprises almost exclusively Annelida. The similar weighted average Δ^13^C values in plantations and rainforest suggest that from an energetic perspective, soil food webs in plantations are as efficient in processing old organic carbon as in rainforest, despite having a very different structure. However, vulnerability of such systems against future changes is questionable since their functioning relies on a single detritivore group comprising predominantly single invasive species (A. Potapov et al., 2021).

Unlike Δ^13^C, changes in Δ^15^N with changes in land use were less consistent across animal groups. Lower Δ^15^N values in Oribatida in rainforest compared to plantations may be linked to their trophic plasticity, allowing them to shift their trophic position from primary decomposer in rainforest to secondary decomposer in litter-poor plantations (Krause et al., 2019). By contrast, many taxa had lower trophic positions in oil palm compared to rainforest, including Hemiptera, Orthoptera, Isoptera and Preudosorpionida. The low Δ^15^N values in these taxa at least in part may be due to feeding on resources depleted in Δ^15^N relative to litter, such as algae and lichens (Chahartaghi et al., 2005; Schneider et al., 2004), suggesting that non-vascular plants may play a more important role for groups such as Hemiptera, Orthoptera and Isoptera in plantations than in rainforest, potentially associated with the more open canopies in plantations.

The range of Δ^15^N values in soil communities reflects the length of food chains (Cabana and Rasmussen, 1994; Scheu and Falca, 2000), and was the largest in jungle rubber. This was caused by the very low Δ^15^N values of Pauropoda (−5.1‰) and Orthoptera (−2.1‰) and high Δ^15^N values of Diplura (10.0‰). Jungle rubber is a system that is highly heterogeneous in management practices and plant richness (Gouyon et al., 1993; Rembold et al., 2017b), with species richness in some arthropod predators even exceeding that in rainforest at our study sites (Junggebauer et al., 2021). Anthropogenic disturbances in jungle rubber are moderate compared to monoculture plantation systems (Barnes et al., 2014) and food-chains have been found to be longest at intermediate levels of disturbance (Menge and Sutherland, 1987; Polis and Winemiller, 2018; David M. Post, 2002), which may explain the largest range of Δ^15^N values in jungle rubber.

### 3. Changes in functional diversity of soil food webs from community and energetic perspectives

Refuting our second hypothesis, neither isotopic diversity nor isotopic redundancy were higher in rainforest than in plantations. However, isotopic richness was slightly higher in the two more natural systems (i.e., rainforest and jungle rubber) than in rubber and oil palm plantations. Oil palm showed significantly higher community dispersion values than the other land-use systems and in trend had the highest community divergence (unweighted values) reflecting the proportion of groups with the most extreme trophic (isotopic) niches within the community (Cucherousset and Villéger, 2015; Mason et al., 2005; Villéger et al., 2008). At least in part this likely was due to feeding on non-vascular plants, such as algae and lichens, characterized by very different stable isotope values than C3 plants, i.e. the dominant vegetation at our study sites (Anton M. Potapov et al., 2019). As discussed above, the more open canopy in plantations favours algae and lichens (Drescher et al., 2016; Schulz et al., 2019), together with the monopolization of detrital channel by earthworms (representing another ‘extreme’ isotopic niche), use of non-vascular plants explains high dispersion and divergence of energy channeling in oil palm plantations.

From the ‘energetic perspective’, soil food web divergence in plantations was significantly higher than in rainforest. This contrasts previous evidence that functional divergence decreases with disturbance (Gerisch et al., 2012; Mouillot et al., 2013). However, contrary to divergence, energetic dispersion was in trend higher in rainforest than in the other land-use systems. Similar to the community metrics, the energetic metrics indicated that food web characteristics in plantations deviate from those in rainforest (high divergence), with food webs being less balanced (low dispersion) with most of the energy being channeled and locked into earthworms.

Community isotopic uniqueness, defined as the inverse of the average isotopic redundancy, and community evenness (Cucherousset and Villéger, 2015), were low in jungle rubber. Most of the soil animal groups clustered in a small region in stable isotope space in jungle rubber, resulting in low community uniqueness, while Pauropoda and Orthoptera were far from this cluster, resulting in low community evenness. High functional redundancy may buffer against land-use impacts and promote stable food webs with long trophic chains (Brodie et al., 2014; Chua et al., 2021; Sanders et al., 2018), which is supported by the results of our study. The increase in isotopic uniqueness (both energetic and unweighted) in oil palm plantations may reflect eroded resilience of this system against future changes.

Overall, community food-web metrics were more sensitive to changes in land use than energetic food-web metrics, suggesting that compositional changes in soil food webs with land use are stronger than changes in energy channeling. This echoes earlier findings that land-use effects on soil animal biodiversity exceed those on functional diversity (Potapov et al., 2020). Results of our SEM indicated significant direct effects of land use-induced environmental changes (i.e., litter amount, tree density and tree species richness) on community food-web metrics. The amount and quality of leaf litter are important drivers of soil fauna composition and soil food-web structure, being both the food and the habitat for soil animals (Fujii et al., 2020; Sayer et al., 2006). Apart from litter-mediated effects, changes in tree density and species richness are associated with changes in root-derived resources (Ballauff et al., 2021), which also fuel belowground food webs (Bradford, 2016; Pollierer et al., 2007), and this may explain the direct effects of tree communities on the food-web metrics. By contrast, energetic food-web metrics were not directly affected by changes in tree communities and pH, but were linked to the changes in earthworms’ abundance. High soil pH favors colonization of plantations by earthworms and this is common in the tropics (Marichal et al., 2010; A. Potapov et al., 2021). The close association between energetic food-web metrics and the fraction earthworms contribute to community metabolism stems to a large extent from the mathematical dependence between these two variables. However, we intentionally wanted to illustrate those strong shifts in the functional diversity of food webs may result from a single group benefiting from certain environmental changes.

In conclusion, our study is among the first comprehensive assessments of tropical soil food webs and their variation due to land-use changes. Low Δ^13^C values in most soil animal groups in plantations in comparison to rainforest indicate a shift towards using plant carbon and ‘fast’ energy channeling based on high quality understory plants (weeds) as well as algae. On the other hand, the trophic niche of earthworms as major macrofauna detritivores stayed unchanged and they monopolized the ‘slow’ detrital channel in plantations. This resulted in systems with strong divergence and imbalance in energetic pathways potentially compromising functional stability of plantation systems. Other studies at our sites showed that these changes in soil food web characteristics with transformation of rainforest into plantations are associated with reduced soil functioning (Grass et al., 2020) and litter invertebrate biodiversity (Barnes et al., 2014). Our analyzes allowed to uncover the mechanisms responsible for these changes and demonstrated that land-use effects on soil biodiversity from a ‘community perspective’ are in part buffered from the perspective of energy channeling (‘energetic perspective’), but resistance of plantations against future changes in climate and land use may be compromised.

## Acknowledgement

This study was funded by the Deutsche Forschungsgemeinschaft (DFG), project number 192626868–SFB 990 in the framework of the collaborative German-Indonesian research project CRC990. Z.Z are supported by China Scholarship Council (CSC) (202004910314). We thank Dr. Katja Rembold and Prof. Holger Kreft for providing vegetation parameters; we also thank Zhijing Xie and Haifeng Yin for discussion. Special gratitude goes to Svenja Meyer for the animal silhouettes.

## References

Ågren GI, Bosatta E, Balesdent J. 1996. Isotope discrimination during decomposition of organic matter: a theoretical analysis. Soil Sci Soc Am J 60:1121–1126. doi:10.2136/sssaj1996.03615995006000040023x

Ballauff J, Schneider D, Edy N, Irawan B, Daniel R, Polle A. 2021. Shifts in root and soil chemistry drive the assembly of belowground fungal communities in tropical land-use systems. Soil Biol Biochem 154:108140. doi:10.1016/j.soilbio.2021.108140

Bardgett RD, van der Putten WH. 2014. Belowground biodiversity and ecosystem functioning. Nature 515:505–511. doi:10.1038/nature13855

Bardgett RD, Wardle DA. 2010. Aboveground-belowground linkages: biotic interactions, ecosystem processes, and global change, Oxford series in ecology and evolution. Oxford: Oxford University Press.

Barnes AD, Jochum M, Lefcheck JS, Eisenhauer N, Scherber C, O’Connor MI, de Ruiter P, Brose U. 2018. Energy Flux: The Link between Multitrophic Biodiversity and Ecosystem Functioning. Trends Ecol Evol 33:186–197. doi:10.1016/j.tree.2017.12.007

Barnes AD, Jochum M, Mumme S, Haneda NF, Farajallah A, Widarto TH, Brose U. 2014. Consequences of tropical land use for multitrophic biodiversity and ecosystem functioning. Nat Commun 5. doi:10.1038/ncomms6351

Blouin M, Hodson ME, Delgado EA, Baker G, Brussaard L, Butt KR, Dai J, Dendooven L, Peres G, Tondoh JE, Cluzeau D, Brun J-J. 2013. A review of earthworm impact on soil function and ecosystem services: Earthworm impact on ecosystem services. Eur J Soil Sci 64:161–182. doi:10.1111/ejss.12025

Bluhm SL, Potapov AM, Shrubovych J, Ammerschubert S, Polle A, Scheu S. 2019. Protura are unique: first evidence of specialized feeding on ectomycorrhizal fungi in soil invertebrates. BMC Ecol 19:10. doi:10.1186/s12898-019-0227-y

Bommarco R, Kleijn D, Potts SG. 2013. Ecological intensification: harnessing ecosystem services for food security. Trends Ecol Evol 28:230–238. doi:10.1016/j.tree.2012.10.012

Boström B, Comstedt D, Ekblad A. 2007. Isotope fractionation and 13C enrichment in soil profiles during the decomposition of soil organic matter. Oecologia 153:89–98. doi:10.1007/s00442-007-0700-8

Bradford MA. 2016. Re-visioning soil food webs. Soil Biol Biochem 102:1–3. doi:10.1016/j.soilbio.2016.08.010

Brodie JF, Aslan CE, Rogers HS, Redford KH, Maron JL, Bronstein JL, Groves CR. 2014. Secondary extinctions of biodiversity. Trends Ecol Evol 29:664–672. doi:10.1016/j.tree.2014.09.012

Brose U, Scheu S. 2014. Into darkness: unravelling the structure of soil food webs. Oikos 123:1153–1156. doi:10.1111/oik.01768

Brown JH, Gillooly JF, Allen AP, Savage VM, West GB. 2004. TOWARD A METABOLIC THEORY OF ECOLOGY. Ecology 85:1771–1789. doi:10.1890/03-9000

Cabana G, Rasmussen JB. 1994. Modeling Food-Chain Structure and Contaminant Bioaccumulation Using Stable Nitrogen Isotopes. Nature 372:255–257. doi:10.1038/372255a0

Chahartaghi M, Langel R, Scheu S, Ruess L. 2005. Feeding guilds in Collembola based on nitrogen stable isotope ratios. Soil Biol Biochem 37:1718–1725. doi:10.1016/j.soilbio.2005.02.006

Chua KWJ, Liew JH, Wilkinson CL, Ahmad AB, Tan HH, Yeo DCJ. 2021. Land-use change erodes trophic redundancy in tropical forest streams: Evidence from amino acid stable isotope analysis. J Anim Ecol 1365–2656.13462. doi:10.1111/1365-2656.13462

Clough Y, Krishna VV, Corre MD, Darras K, Denmead LH, Meijide A, Moser S, Musshoff O, Steinebach S, Veldkamp E, Allen K, Barnes AD, Breidenbach N, Brose U, Buchori D, Daniel R, Finkeldey R, Harahap I, Hertel D, Holtkamp AM, Horandl E, Irawan B, Jaya NS, Jochum M, Klarner B, Knohl A, Kotowska MM, Krashevska V, Kreft H, Kurniawan S, Leuschner C, Maraun M, Melati DN, Opfermann N, Perez-Cruzado C, Prabowo WE, Rembold K, Rizali A, Rubiana R, Schneider D, Tjitrosoedirdjo SS, Tjoa A, Tscharntke T, Scheu S. 2016. Land-use choices follow profitability at the expense of ecological functions in Indonesian smallholder landscapes. Nat Commun 7. doi:10.1038/ncomms13137

Coleman DC, Reid CPP, Cole CV. 1983. Biological Strategies of Nutrient Cycling in Soil SystemsAdvances in Ecological Research. Elsevier. pp. 1–55. doi:10.1016/S0065-2504(08)60107-5

Cucherousset J, Villéger S. 2015. Quantifying the multiple facets of isotopic diversity: New metrics for stable isotope ecology. Ecol Indic 56:152–160. doi:10.1016/j.ecolind.2015.03.032

Darras KFA, Corre MD, Formaglio G, Tjoa A, Potapov A, Brambach F, Sibhatu KT, Grass I, Rubiano AA, Buchori D, Drescher J, Fardiansah R, Hölscher D, Irawan B, Kneib T, Krashevska V, Krause A, Kreft H, Li K, Maraun M, Polle A, Ryadin AR, Rembold K, Stiegler C, Scheu S, Tarigan S, Valdés-Uribe A, Yadi S, Tscharntke T, Veldkamp E. 2019. Reducing Fertilizer and Avoiding Herbicides in Oil Palm Plantations—Ecological and Economic Valuations. Front For Glob Change 2:65. doi:10.3389/ffgc.2019.00065

de Vries FT, Hoffland E, van Eekeren N, Brussaard L, Bloem J. 2006. Fungal/bacterial ratios in grasslands with contrasting nitrogen management. Soil Biol Biochem 38:2092–2103. doi:10.1016/j.soilbio.2006.01.008

de Vries FT, Liiri ME, Bjørnlund L, Bowker MA, Christensen S, Setälä HM, Bardgett RD. 2012. Land use alters the resistance and resilience of soil food webs to drought. Nat Clim Change 2:276–280. doi:10.1038/nclimate1368

de Vries FT, Thebault E, Liiri M, Birkhofer K, Tsiafouli MA, Bjornlund L, Bracht Jorgensen H, Brady MV, Christensen S, de Ruiter PC, d’Hertefeldt T, Frouz J, Hedlund K, Hemerik L, Hol WHG, Hotes S, Mortimer SR, Setala H, Sgardelis SP, Uteseny K, van der Putten WH, Wolters V, Bardgett RD. 2013. Soil food web properties explain ecosystem services across European land use systems. Proc Natl Acad Sci 110:14296–14301. doi:10.1073/pnas.1305198110

Dobrovolski R, Diniz-Filho JAF, Loyola RD, De Marco Júnior P. 2011. Agricultural expansion and the fate of global conservation priorities. Biodivers Conserv 20:2445–2459. doi:10.1007/s10531-011-9997-z

Drescher J, Rembold K, Allen K, Beckschafer P, Buchori D, Clough Y, Faust H, Fauzi AM, Gunawan D, Hertel D, Irawan B, Jaya INS, Klarner B, Kleinn C, Knohl A, Kotowska MM, Krashevska V, Krishna V, Leuschner C, Lorenz W, Meijide A, Melati D, Nomura M, Perez-Cruzado C, Qaim M, Siregar IZ, Steinebach S, Tjoa A, Tscharntke T, Wick B, Wiegand K, Kreft H, Scheu S. 2016. Ecological and socio-economic functions across tropical land use systems after rainforest conversion. Philos Trans R Soc B-Biol Sci 371. doi:10.1098/rstb.2015.0275

Ehnes RB, Rall BC, Brose U. 2011. Phylogenetic grouping, curvature and metabolic scaling in terrestrial invertebrates: Invertebrate metabolism. Ecol Lett 14:993–1000. doi:10.1111/j.1461-0248.2011.01660.x

Fujii S, Berg MP, Cornelissen JHC. 2020. Living Litter: Dynamic Trait Spectra Predict Fauna Composition. Trends Ecol Evol 35:886–896. doi:10.1016/j.tree.2020.05.007

Fujii S, Haraguchi TF, Tayasu I. 2021. Radiocarbon signature reveals that most springtails depend on carbon from living plants. Biol Lett 17:20210353. doi:10.1098/rsbl.2021.0353

Geisen S, Briones MJI, Gan H, Behan-Pelletier VM, Friman V-P, de Groot GA, Hannula SE, Lindo Z, Philippot L, Tiunov AV, Wall DH. 2019. A methodological framework to embrace soil biodiversity. Soil Biol Biochem 136:107536. doi:10.1016/j.soilbio.2019.107536

Gerisch M, Agostinelli V, Henle K, Dziock F. 2012. More species, but all do the same: contrasting effects of flood disturbance on ground beetle functional and species diversity. Oikos 121:508–515. doi:10.1111/j.1600-0706.2011.19749.x

Gessner MO, Swan CM, Dang CK, McKie BG, Bardgett RD, Wall DH, Hättenschwiler S. 2010. Diversity meets decomposition. Trends Ecol Evol 25:372–380. doi:10.1016/j.tree.2010.01.010

Gouyon A, de Foresta H, Levang P. 1993. Does ‘jungle rubber’ deserve its name? An analysis of rubber agroforestry systems in southeast Sumatra. Agrofor Syst 22:181–206. doi:10.1007/BF00705233

Grass I, Kubitza C, Krishna VV, Corre MD, Mußhoff O, Pütz P, Drescher J, Rembold K, Ariyanti ES, Barnes AD, Brinkmann N, Brose U, Brümmer B, Buchori D, Daniel R, Darras KFA, Faust H, Fehrmann L, Hein J, Hennings N, Hidayat P, Hölscher D, Jochum M, Knohl A, Kotowska MM, Krashevska V, Kreft H, Leuschner C, Lobite NJS, Panjaitan R, Polle A, Potapov AM, Purnama E, Qaim M, Röll A, Scheu S, Schneider D, Tjoa A, Tscharntke T, Veldkamp E, Wollni M. 2020. Trade-offs between multifunctionality and profit in tropical smallholder landscapes. Nat Commun 11:1186. doi:10.1038/s41467-020-15013-5

Guerra CA, Bardgett RD, Caon L, Crowther TW, Delgado-Baquerizo M, Montanarella L, Navarro LM, Orgiazzi A, Singh BK, Tedersoo L, Vargas-Rojas R, Briones MJI, Buscot F, Cameron EK, Cesarz S, Chatzinotas A, Cowan DA, Djukic I, van den Hoogen J, Lehmann A, Maestre FT, Marín C, Reitz T, Rillig MC, Smith LC, de Vries FT, Weigelt A, Wall DH, Eisenhauer N. 2021. Tracking, targeting, and conserving soil biodiversity. Science 371:239–241. doi:10.1126/science.abd7926

Guillaume T, Kotowska MM, Hertel D, Knohl A, Krashevska V, Murtilaksono K, Scheu S, Kuzyakov Y. 2018. Carbon costs and benefits of Indonesian rainforest conversion to plantations. Nat Commun 9:2388. doi:10.1038/s41467-018-04755-y

Hooper DU, Bignell DE, Brown VK, Brussard L, Mark Dangerfield J, Wall DH, Wardle DA, Coleman DC, Giller KE, Lavelle P, Van Der Putten WH, De Ruiter PC, Rusek J, Silver WL, Tiedje JM, Wolters V. 2000. Interactions between Aboveground and Belowground Biodiversity in Terrestrial Ecosystems: Patterns, Mechanisms, and Feedbacks. BioScience 50:1049. doi:10.1641/0006-3568(2000)050[1049:IBAABB]2.0.CO;2

Hunt HW, Coleman DC, Ingham ER, Ingham RE, Elliott ET, Moore JC, Rose SL, Reid CPP, Morley CR. 1987. The detrital food web in a shortgrass prairie. Biol Fertil Soils 3–3. doi:10.1007/BF00260580

Hyodo F. 2015. Use of stable carbon and nitrogen isotopes in insect trophic ecology: Use of isotope in insect trophic ecology. Entomol Sci 18:295–312. doi:10.1111/ens.12128

Hyodo F, Matsumoto T, Takematsu Y, Kamoi T, Fukuda D, Nakagawa M, Itioka T. 2010. The structure of a food web in a tropical rain forest in Malaysia based on carbon and nitrogen stable isotope ratios. J Trop Ecol 26:205–214. doi:10.1017/S0266467409990502

Hyodo F, Uchida T, Kaneko N, Tayasu I. 2012. Use of radiocarbon to estimate diet ages of earthworms across different climate regions. Appl Soil Ecol 62:178–183. doi:10.1016/j.apsoil.2012.09.014

Illig J, Langel R, Norton RA, Scheu S, Maraun M. 2005. Where are the decomposers? Uncovering the soil food web of a tropical montane rain forest in southern Ecuador using stable isotopes ( ^15^N). J Trop Ecol 21:589–593. doi:10.1017/S0266467405002646

Junggebauer A, Hartke TR, Ramos D, Schaefer I, Buchori D, Hidayat P, Scheu S, Drescher J. 2021. Changes in diversity and community assembly of jumping spiders (Araneae: Salticidae) after rainforest conversion to rubber and oil palm plantations. PeerJ 9:e11012. doi:10.7717/peerj.11012

Kempson D, Lloyd M, Ghelardi R. 1963. A new extractor for woodland litter. Pedobiologia 3:1–21.

Klarner B, Winkelmann H, Krashevska V, Maraun M, Widyastuti R, Scheu S. 2017. Trophic niches, diversity and community composition of invertebrate top predators (Chilopoda) as affected by conversion of tropical lowland rainforest in Sumatra (Indonesia). PLOS ONE 12:e0180915. doi:10.1371/journal.pone.0180915

Koh LP, Ghazoul J. 2010. Spatially explicit scenario analysis for reconciling agricultural expansion, forest protection, and carbon conservation in Indonesia. Proc Natl Acad Sci 107:11140–11144. doi:10.1073/pnas.1000530107

Kotowska MM, Leuschner C, Triadiati T, Meriem S, Hertel D. 2015. Quantifying above- and belowground biomass carbon loss with forest conversion in tropical lowlands of Sumatra (Indonesia). Glob Change Biol 21:3620–3634. doi:10.1111/gcb.12979

Krashevska V, Klarner B, Widyastuti R, Maraun M, Scheu S. 2015. Impact of tropical lowland rainforest conversion into rubber and oil palm plantations on soil microbial communities. Biol Fertil Soils 51:697–705. doi:10.1007/s00374-015-1021-4

Krashevska V, Malysheva E, Klarner B, Mazei Y, Maraun M, Widyastuti R, Scheu S. 2018. Micro-decomposer communities and decomposition processes in tropical lowlands as affected by land use and litter type. Oecologia 187:255–266. doi:10.1007/s00442-018-4103-9

Krashevska V, Sandmann D, Marian F, Maraun M, Scheu S. 2017. Leaf Litter Chemistry Drives the Structure and Composition of Soil Testate Amoeba Communities in a Tropical Montane Rainforest of the Ecuadorian Andes. Microb Ecol 74:681–690. doi:10.1007/s00248-017-0980-4

Krause A, Sandmann D, Bluhm SL, Ermilov S, Widyastuti R, Haneda NF, Scheu S, Maraun M. 2019. Shift in trophic niches of soil microarthropods with conversion of tropical rainforest into plantations as indicated by stable isotopes (15N, 13C). PLOS ONE 14:e0224520. doi:10.1371/journal.pone.0224520

Krause A, Sandmann D, Potapov A, Ermilov S, Widyastuti R, Haneda NF, Scheu S, Maraun M. 2021. Variation in community-level trophic niches of soil microarthropods with conversion of tropical rainforest into plantation systems as indicated by stable isotopes (15N, 13C). Front Ecol Evol 9:592149. doi:10.3389/fevo.2021.592149

Laliberté E, Legendre P. 2010. A distance-based framework for measuring functional diversity from multiple traits. Ecology 91:299–305. doi:10.1890/08-2244.1

Laurance WF. 2007. Have we overstated the tropical biodiversity crisis? Trends Ecol Evol 22:65–70. doi:10.1016/j.tree.2006.09.014

Layman CA, Araujo MS, Boucek R, Hammerschlag-Peyer CM, Harrison E, Jud ZR, Matich P, Rosenblatt AE, Vaudo JJ, Yeager LA, Post DM, Bearhop S. 2012. Applying stable isotopes to examine food-web structure: an overview of analytical tools. Biol Rev 87:545–562. doi:10.1111/j.1469-185X.2011.00208.x

Layman CA, Arrington DA, Montana CG, Post DM. 2007. Can stable isotope ratios provide for community-wide measures of trophic structure? Ecology 88:42–48. doi:10.1890/0012-9658(2007)88[42:Csirpf]2.0.Co;2

Liebke DF, Harms D, Widyastuti R, Scheu S, Potapov AM. 2021. Impact of rainforest conversion into monoculture plantation systems on pseudoscorpion density, diversity and trophic niches. doi:10.25674/SO93ISS2ID147

Margono BA, Turubanova S, Zhuravleva I, Potapov P, Tyukavina A, Baccini A, Goetz S, Hansen MC. 2012. Mapping and monitoring deforestation and forest degradation in Sumatra (Indonesia) using Landsat time series data sets from 1990 to 2010. Environ Res Lett 7:034010. doi:10.1088/1748-9326/7/3/034010

Marichal R, Martinez AF, Praxedes C, Ruiz D, Carvajal AF, Oszwald J, del Pilar Hurtado M, Brown GG, Grimaldi M, Desjardins T, Sarrazin M, Decaëns T, Velasquez E, Lavelle P. 2010. Invasion of Pontoscolex corethrurus (Glossoscolecidae, Oligochaeta) in landscapes of the Amazonian deforestation arc. Appl Soil Ecol 46:443–449. doi:10.1016/j.apsoil.2010.09.001

Mason NWH, Mouillot D, Lee WG, Wilson JB. 2005. Functional richness, functional evenness and functional divergence: the primary components of functional diversity. Oikos 111:112–118. doi:10.1111/j.0030-1299.2005.13886.x

Matson PA. 1997. Agricultural Intensification and Ecosystem Properties. Science 277:504–509. doi:10.1126/science.277.5325.504

McCann KS, Hastings AG, Huxel GR. 1998. Weak Trophic Interactions and the Balance of Nature. Nature 395:794–798.

McGrath DA, Smith CK, Gholz HL, Oliveira F de A. 2001. Effects of Land-Use Change on Soil Nutrient Dynamics in Amazônia. Ecosystems 4:625–645. doi:10.1007/s10021-001-0033-0

Mendiburu F de. 2020. agricolae: Statistical Procedures for Agricultural Research.

Menge BA, Sutherland JP. 1987. Community Regulation: Variation in Disturbance, Competition, and Predation in Relation to Environmental Stress and Recruitment. Am Nat 130:730–757. doi:10.1086/284741

Miettinen J, Shi C, Liew SC. 2011. Deforestation rates in insular Southeast Asia between 2000 and 2010: DEFORESTATION IN INSULAR SOUTHEAST ASIA 2000-2010. Glob Change Biol 17:2261–2270. doi:10.1111/j.1365-2486.2011.02398.x

Moore JC, McCann K, de Ruiter PC. 2005. Modeling trophic pathways, nutrient cycling, and dynamic stability in soils. Pedobiologia 49:499–510. doi:10.1016/j.pedobi.2005.05.008

Mouillot D, Graham NAJ, Villéger S, Mason NWH, Bellwood DR. 2013. A functional approach reveals community responses to disturbances. Trends Ecol Evol 28:167–177. doi:10.1016/j.tree.2012.10.004

Newbold T, Hudson LN, Hill SLL, Contu S, Lysenko I, Senior RA, Börger L, Bennett DJ, Choimes A, Collen B, Day J, De Palma A, Díaz S, Echeverria-Londoáo S, Edgar MJ, Feldman A, Garon M, Harrison MLK, Alhusseini T, Ingram DJ, Itescu Y, Kattge J, Kemp V, Kirkpatrick L, Kleyer M, Correia DLP, Martin CD, Meiri S, Novosolov M, Pan Y, Phillips HRP, Purves DW, Robinson A, Simpson J, Tuck SL, Weiher E, White HJ, Ewers RM, Mace GM, Scharlemann JPW, Purvis A. 2015. Global effects of land use on local terrestrial biodiversity. Nature 520:45–50. doi:10.1038/nature14324

Parnell AC, Inger R, Bearhop S, Jackson AL. 2010. Source Partitioning Using Stable Isotopes: Coping with Too Much Variation. Plos One 5. doi:10.1371/journal.pone.0009672

Petchey OL, Gaston KJ. 2006. Functional diversity: back to basics and looking forward. Ecol Lett 9:741–758. doi:10.1111/j.1461-0248.2006.00924.x

Peterson BJ, Fry B. 1987. Stable Isotopes in Ecosystem Studies. Annu Rev Ecol Syst 18:293–320. doi:10.1146/annurev.es.18.110187.001453

Polis GA, Winemiller KO, editors. 2018. Food webs: integration of patterns & dynamics, Softcover reprint of the harcover 1st edition 1996. ed. Dordrecht: Springer-Science+Business Media, B.V.

Pollierer MM, Langel R, Körner C, Maraun M, Scheu S. 2007. The underestimated importance of belowground carbon input for forest soil animal food webs. Ecol Lett 10:729–736. doi:10.1111/j.1461-0248.2007.01064.x

Pollierer MM, Langel R, Scheu S, Maraun M. 2009. Compartmentalization of the soil animal food web as indicated by dual analysis of stable isotope ratios (15N/14N and 13C/12C). Soil Biol Biochem 41:1221–1226. doi:10.1016/j.soilbio.2009.03.002

Post D.M. 2002. Using stable isotopes to estimate trophic position: Models, methods, and assumptions. Ecology 83:703–718. doi:10.1890/0012-9658(2002)083[0703:Usitet]2.0.Co;2

Post David M. 2002. The long and short of food-chain length. Trends Ecol Evol 17:269–277. doi:10.1016/S0169-5347(02)02455-2

Potapov A, Schaefer I, Jochum M, Widyastuti R, Eisenhauer N, Scheu S. 2021. Oil palm and rubber expansion facilitates earthworm invasion in Indonesia. Biol Invasions. doi:10.1007/s10530-021-02539-y

Potapov AM, Dupérré N, Jochum M, Dreczko K, Klarner B, Barnes AD, Krashevska V, Rembold K, Kreft H, Brose U, Widyastuti R, Harms D, Scheu S. 2020. Functional losses in ground spider communities due to habitat structure degradation under tropical land-use change. Ecology 101. doi:10.1002/ecy.2957

Potapov A. M., Klarner B, Sandmann D, Widyastuti R, Scheu S. 2019a. Linking size spectrum, energy flux and trophic multifunctionality in soil food webs of tropical land-use systems. J Anim Ecol 88:1845–1859. doi:10.1111/1365-2656.13027

Potapov AM, Korotkevich AYu, Tiunov AV. 2018. Non-vascular plants as a food source for litter-dwelling Collembola: Field evidence. Pedobiologia 66:11–17. doi:10.1016/j.pedobi.2017.12.005

Potapov AM, Rozanova OL, Semenina EE, Leonov VD, Belyakova OI, Bogatyreva VYu, Degtyarev MI, Esaulov AS, Korotkevich AYu, Kudrin AA, Malysheva EA, Mazei YA, Tsurikov SM, Zuev AG, Tiunov AV. 2021. Size compartmentalization of energy channeling in terrestrial belowground food webs. Ecology. doi:10.1002/ecy.3421

Potapov A. M., Scheu S, Tiunov AV. 2019b. Trophic consistency of supraspecific taxa in below-ground invertebrate communities: Comparison across lineages and taxonomic ranks. Funct Ecol 33:1172–1183. doi:10.1111/1365-2435.13309

Potapov AM, Semenina EE, Kurakov AV, Tiunov AV. 2013. Large 13C/12C and small 15N/14N isotope fractionation in an experimental detrital foodweb (litter–fungi–collembolans). Ecol Res 28:1069–1079. doi:10.1007/s11284-013-1088-z

Potapov Anton M., Tiunov AV, Scheu S. 2019. Uncovering trophic positions and food resources of soil animals using bulk natural stable isotope composition. Biol Rev 94:37–59. doi:10.1111/brv.12434

R Core Team. 2020. R: A Language and Environment for Statistical Computing. Vienna, Austria: R Foundation for Statistical Computing.

Rembold K, Mangopo H, Tjitrosoedirdjo SS, Kreft H. 2017a. Plant diversity, forest dependency, and alien plant invasions in tropical agricultural landscapes. Biol Conserv 213:234–242. doi:10.1016/j.biocon.2017.07.020

Rembold K, Tjitrosoedirdjo SS, Kreft H. 2017b. Common wayside plants of Jambi Province (Sumatra, Indonesia). Biodiversity, Macroecology and Biogeography, Faculty of Forest Sciences and Forest Ecology of the University of Goettingen. doi:10.3249/WEBDOC-3979

Rooney N, McCann K, Gellner G, Moore JC. 2006. Structural asymmetry and the stability of diverse food webs. Nature 442:265–269. doi:10.1038/nature04887

Rooney N, McCann KS. 2012. Integrating food web diversity, structure and stability. Trends Ecol Evol 27:40–46. doi:10.1016/j.tree.2011.09.001

Rosseel Y. 2012. lavaan: An R Package for Structural Equation Modeling. J Stat Softw 48:1–36.

RStudio Team. 2020. RStudio: Integrated Development Environment for R. Boston, MA: RStudio, PBC.

Sanders D, Thébault E, Kehoe R, Frank van Veen FJ. 2018. Trophic redundancy reduces vulnerability to extinction cascades. Proc Natl Acad Sci 115:2419–2424. doi:10.1073/pnas.1716825115

Sayer EJ, Tanner EVJ, Lacey AL. 2006. Effects of litter manipulation on early-stage decomposition and meso-arthropod abundance in a tropical moist forest. For Ecol Manag 229:285–293. doi:https://doi.org/10.1016/j.foreco.2006.04.007

Schermelleh-Engel K, Moosbrugger H, Müller H. 2003. Evaluating the Fit of Structural Equation Models: Tests of Significance and Descriptive Goodness-of-Fit Measures 8:53.

Scheu S, Falca M. 2000. The soil food web of two beech forests (Fagus sylvatica) of contrasting humus type: stable isotope analysis of a macro- and a mesofauna-dominated community. Oecologia 123:285–296. doi:10.1007/s004420051015

Schmitz OJ, Leroux SJ. 2020. Food Webs and Ecosystems: Linking Species Interactions to the Carbon Cycle. Annu Rev Ecol Evol Syst 51:271–295. doi:10.1146/annurev-ecolsys-011720-104730

Schneider K, Migge S, Norton RA, Scheu S, Langel R, Reineking A, Maraun M. 2004. Trophic niche differentiation in soil microarthropods (Oribatida, Acari): evidence from stable isotope ratios (15N/14N). Soil Biol Biochem 36:1769–1774. doi:10.1016/j.soilbio.2004.04.033

Schulz G, Schneider D, Brinkmann N, Edy N, Daniel R, Polle A, Scheu S, Krashevska V. 2019. Changes in trophic groups of protists with conversion of rainforest into rubber and oil palm plantations. Front Microbiol 10:240. doi:10.3389/fmicb.2019.00240

Schwarzmüller F, Eisenhauer N, Brose U. 2015. ‘Trophic whales’ as biotic buffers: weak interactions stabilize ecosystems against nutrient enrichment. J Anim Ecol 84:680–691. doi:10.1111/1365-2656.12324

Sendra A, Jiménez-Valverde A, Selfa J. 2021. Diversity, ecology, distribution and biogeography of Diplura. Insect Conserv Divers 11.

Starling JH. 1944. Ecological Studies of the Pauropoda of the Duke Forest. Ecol Monogr 14:291–310. doi:10.2307/1948445

Steffan SA, Chikaraishi Y, Currie CR, Horn H, Gaines-Day HR, Pauli JN, Zalapa JE, Ohkouchi N. 2015. Microbes are trophic analogs of animals. Proc Natl Acad Sci 112:15119–15124. doi:10.1073/pnas.1508782112

Susanti WI, Pollierer MM, Widyastuti R, Scheu S, Potapov A. 2019. Conversion of rainforest to oil palm and rubber plantations alters energy channels in soil food webs. Ecol Evol 9:9027–9039. doi:10.1002/ece3.5449

Susanti WI, Widyastuti R, Scheu S, Potapov A. 2021. Trophic niche differentiation and utilisation of food resources in Collembola is altered by rainforest conversion to plantation systems. PeerJ 9:e10971. doi:10.7717/peerj.10971

Tiunov AV, Semenina EE, Aleksandrova AV, Tsurikov SM, Anichkin AE, Novozhilov YK. 2015. Stable isotope composition (δ^13^ C and δ^15^ N values) of slime molds: placing bacterivorous soil protozoans in the food web context: Isotopic composition of slime molds. Rapid Commun Mass Spectrom 29:1465–1472. doi:10.1002/rcm.7238

Tsiafouli MA, Thébault E, Sgardelis SP, de Ruiter PC, van der Putten WH, Birkhofer K, Hemerik L, de Vries FT, Bardgett RD, Brady MV, Bjornlund L, Jørgensen HB, Christensen S, Hertefeldt TD, Hotes S, Gera Hol WH, Frouz J, Liiri M, Mortimer SR, Setälä H, Tzanopoulos J, Uteseny K, Pižl V, Stary J, Wolters V, Hedlund K. 2015. Intensive agriculture reduces soil biodiversity across Europe. Glob Change Biol 21:973–985. doi:10.1111/gcb.12752

Tsurikov SM, Goncharov AA, Tiunov AV. 2015. Intra-body variation and ontogenetic changes in the isotopic composition (13C/12C and 15N/14N) of beetles (Coleoptera). Entomol Rev 95:326–333. doi:10.1134/S0013873815030057

Villéger S, Mason NWH, Mouillot D. 2008. NEW MULTIDIMENSIONAL FUNCTIONAL DIVERSITY INDICES FOR A MULTIFACETED FRAMEWORK IN FUNCTIONAL ECOLOGY. Ecology 89:2290–2301. doi:10.1890/07-1206.1

Wickham H. 2016. ggplot2: Elegant Graphics for Data Analysis. Springer-Verlag New York.

Wilkinson CL, Chua KWJ, Fiala R, Liew JH, Kemp V, Hadi Fikri A, Ewers RM, Kratina P, Yeo DCJ. 2021. Forest conversion to oil palm compresses food chain length in tropical streams. Ecology 102. doi:10.1002/ecy.3199

Yang G, Wagg C, Veresoglou SD, Hempel S, Rillig MC. 2018. How Soil Biota Drive Ecosystem Stability. Trends Plant Sci 23:1057–1067. doi:10.1016/j.tplants.2018.09.007

